## Supplementary material for "Deciphering Neuronal Deficit and Protein Profile Changes in Human Brain Organoids from Patients with Creatine Transporter Deficiency": supplemantal figures

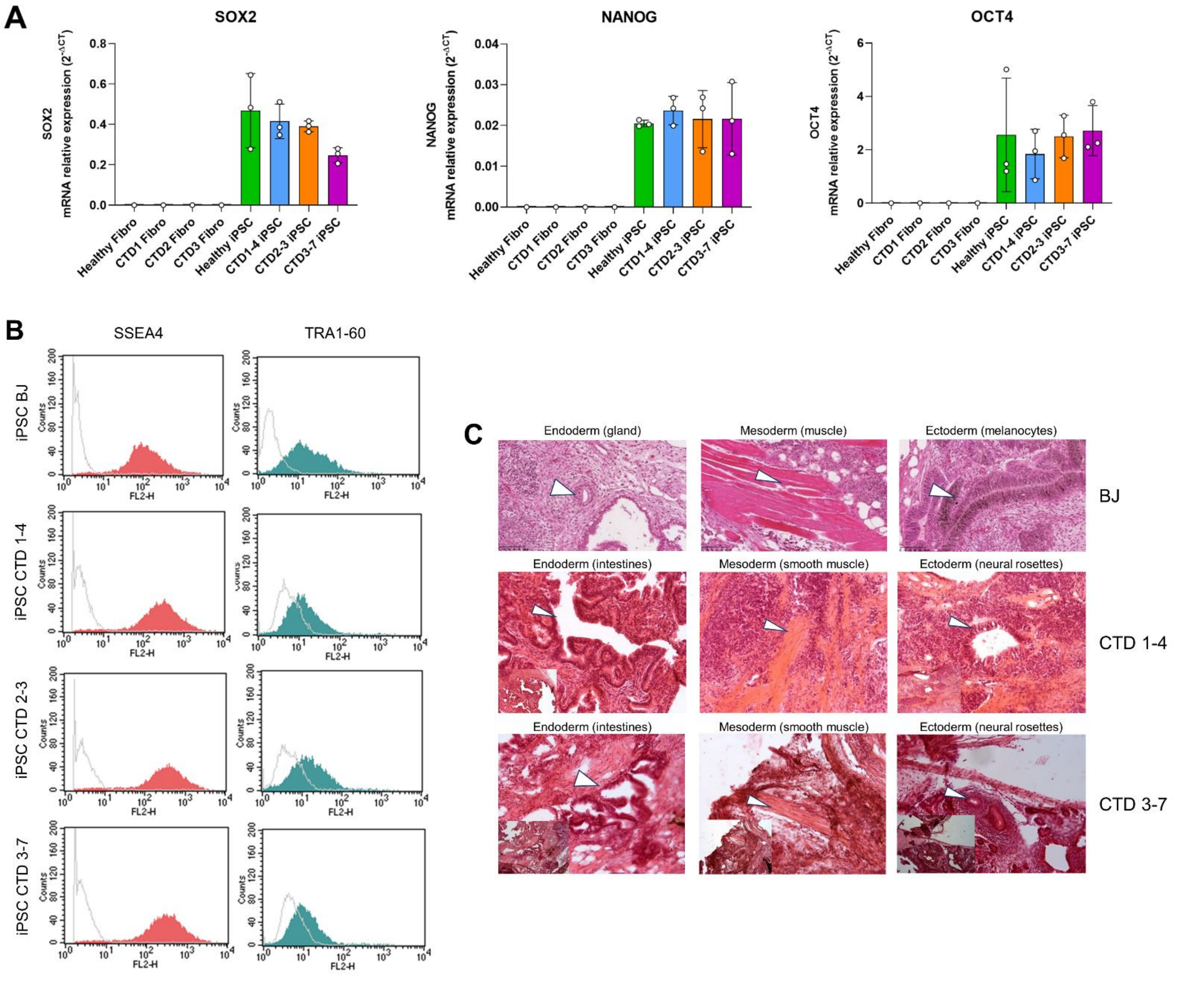


**Figure S1. Generation of CTD iPSCs**

A. RT-qPCR of pluripotency markers SOX2, NANOG and OCT4 in fibroblasts (PK and CTD1, 2, and 3) and iPSCs (BJ and CTD1-4, 2-3, and 3-7). n=1 to 3

B. Representative analysis of SSEA4 and TRA1-60 expression for each iPSC line by flow cytometry. SSEA4 is compared with the appropriate mouse IgG3 control, and TRA1-60 is compared with unlabeled cells. Control: grey line; test: red and green line.

C. Generation of all three germ cell layer components within teratomas from each iPSC line (endoderm, mesoderm, ectoderm). Whole sections of teratoma were performed 5 to 10 weeks of growth and stained with hematoxylin and eosin. Scale bar = 50 µm.

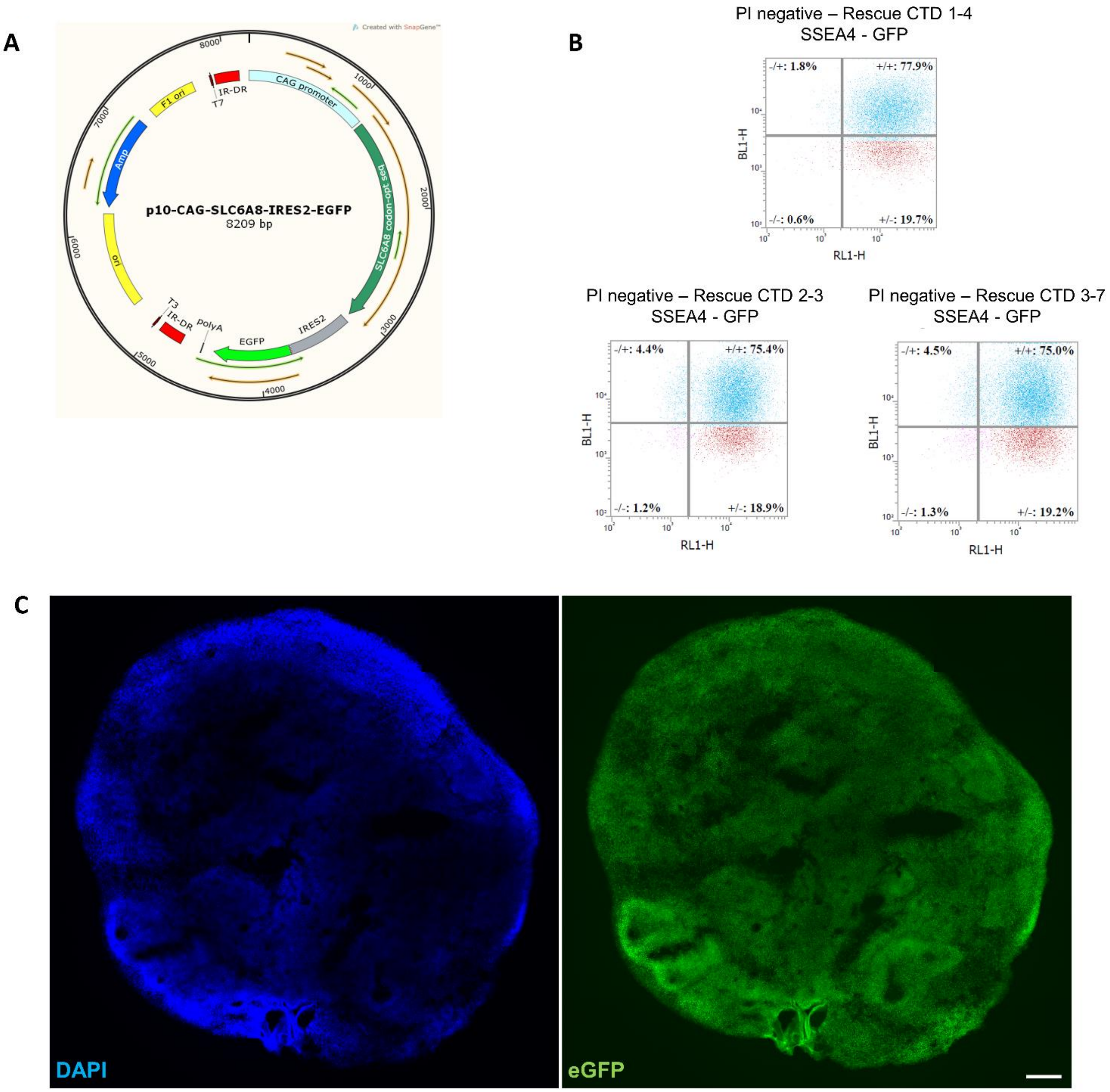


**Figure S2. Generation of CTD-rescue iPSCs**

A. The p10-CAG-SLC6A8-IRES2-eGFP vector used for transfection

B. FACS analysis of transfected cells according to pluripotency marker SSEA4 (RL1-H) and GFP (BL1-H).

C. Representative image of a 2 months old brain organoid tissue section from CTD rescue cell line showing endogenous eGFP fluorescence. DAPI marks nuclei in blue. Scale bar: 200 µm

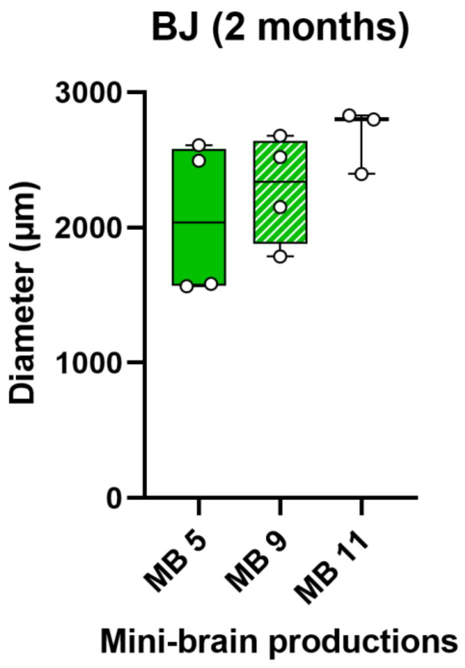


**Figure S3. Assessment of brain organoids variability**

Diameter of healthy BJ brain organoids obtained from 3 different productions.

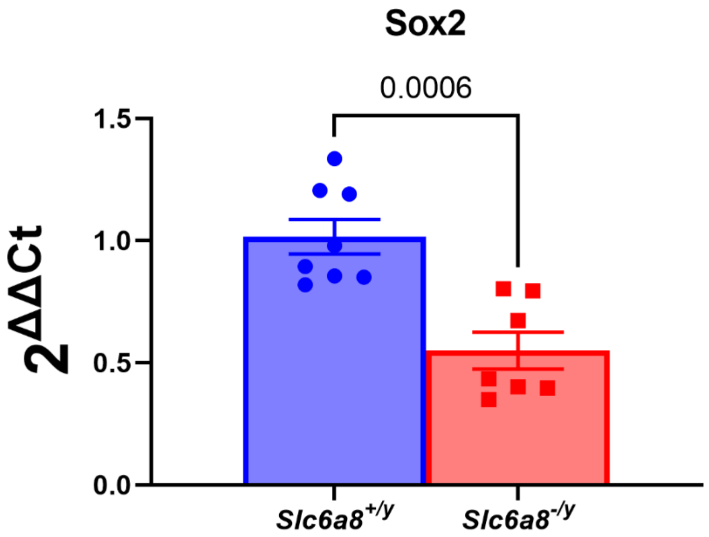


**Figure S4. Neurogenesis deficit in a CTD mouse model**

Relative mRNA expression of SOX2 (radial glial cell markers) in whole brains of postnatal day 0 *Slc6a8*$^{-/y}$ mice. n=8 *Slc6a8*$^{+/y}$, 7 *Slc6a8*$^{-/y}$ mice. Data analyzed using two-tailed t-test on $2^{-(\Delta\Delta Ct)}$ values with mean ΔCt of *Slc6a8*$^{+/y}$ mice used as the control value. t(13)=4.527, P=0.0006

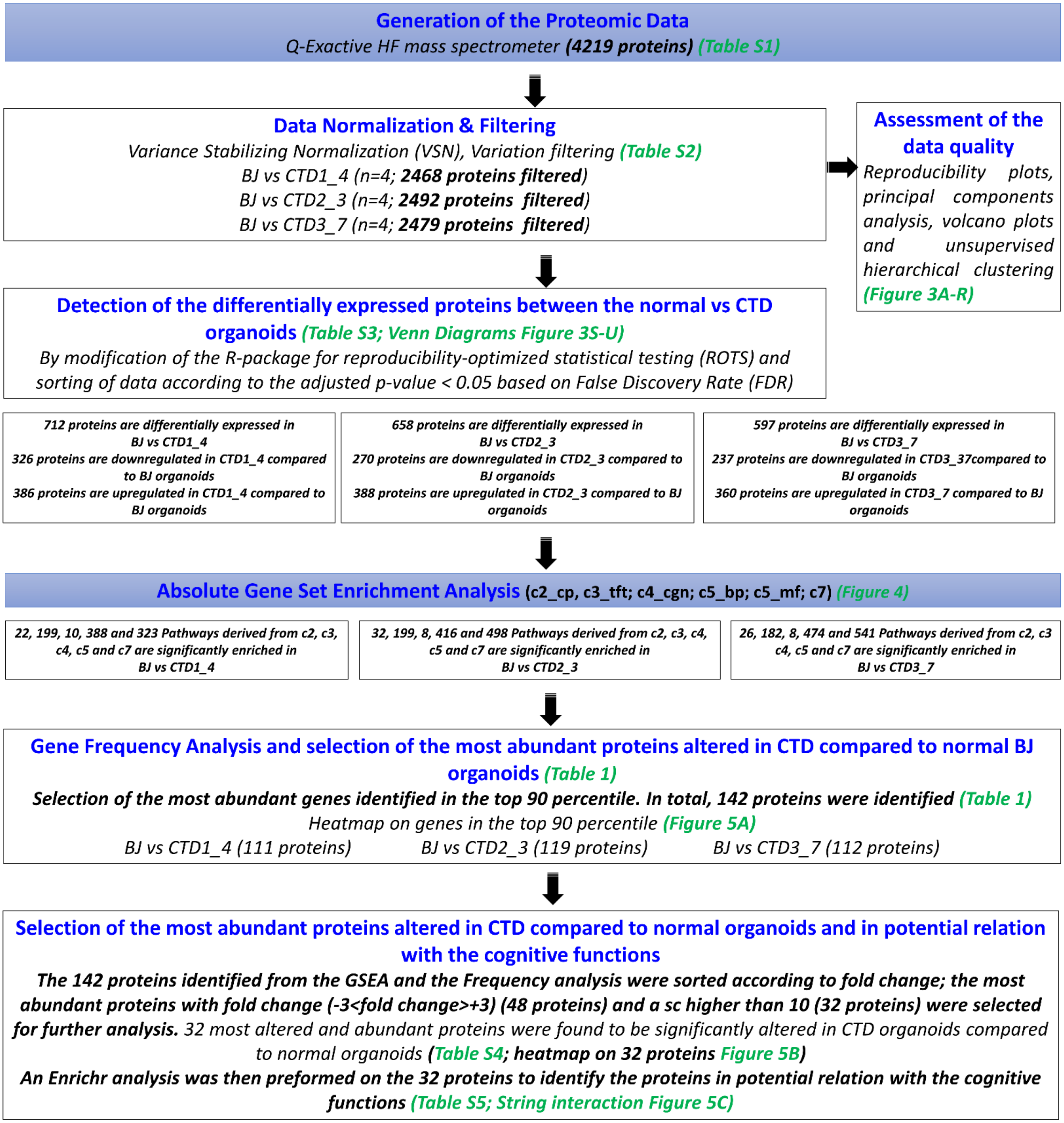


**Figure S5. Flowchart diagram depicting the entire bioinformatics analysis workflow.** The raw data generated by MS/MS was normalized and filtered. The proteins significantly altered in the three CTD-derived cerebral organoids in comparison with healthy cerebral organoids were identified using modification of the R package for ROTS. Absolute GSEA was performed in order to identify the enriched pathways in CTD-derived brain organoids in comparison with healthy cerebral organoids and the list of enriched genes was identified. Their recurrence or frequency in other pathways among all studied gene sets was searched. The frequency analysis identified 142 differentially expressed proteins occurring frequently across all enriched pathways. The 142 proteins identified were then subjected to a disease-related pathway analysis carried out using Enrichr followed by a frequency analysis in order to identify potential functions of the differentially expressed proteins. To further reduce the set of available proteins, the 142 proteins identified from the GSEA and the Frequency analysis were then sorted according to fold change; the most abundant proteins with fold change (-3<fold change>+3) and a sc higher than 10 were retained. 32 most altered and abundant proteins were found to be significantly altered in CTD-derived cerebral organoids compared to normal organoids.
