## supplemental Tables for "Deciphering Neuronal Deficit and Protein Profile Changes in Human Brain Organoids from Patients with Creatine Transporter Deficiency": Table S6 - 32 proteins functions.docx

| **Table S6. 32 proteins functions and relation to CTD symptoms.** | | |
| --- | --- | --- |
| **Protein** | **Functions** | **Potential relation to CTD symptoms or other intellectual disabilities (ID)** |
| RUFY3 | Axon growth | Axon degeneration (Hertz *et al*, 2019) |
| MAP1B | Microtubule-associated protein | Mutations cause ID (Walters *et al*, 2018) FMRP (linked to Fragile X syndrome) (Lu *et al*, 2004) Seizures (Arya *et al*, 2021; Fischer *et al*, 1995) |
| GSK3B | Protein kinase | Rett syndrome (Jorge-Torres *et al*, 2018) Neurodevelopmental disorders with seizures and ID (Fuchs *et al*, 2014, 2015) Down syndrome and neurogenesis (Trazzi *et al*, 2014) Down syndrome and mitochondria defects (Shukkur *et al*, 2006) Fragile X syndrome (Guo *et al*, 2012; Chen *et al*, 2013; Luo *et al*, 2010) Alzeimer's disease (Shaw & Chang, 2013) Epilepsy (Cheng *et al*, 2020) |
| PAK1 | Serine/threonine-protein kinase | ID and seizures (Horn *et al*, 2019; Kernohan *et al*, 2019; Ohori *et al*, 2020) Down syndrome and neurogenesis (Tang *et al*, 2021) Down syndrome and mitochondria (Xu *et al*, 2022) Proliferation of neural progenitor cells (Pan *et al*, 2015) |
| STMN1 | Microtubule regulation | Alzeimer's disease and Down syndrome (Cheon *et al*, 2001) Epilepsy (Zhao *et al*, 2012) |
| FYN | Tyrosine-protein kinase | Linked to tau protein and seizures (Putra *et al*, 2020) |
| GNAI2 | G(i) protein subunit | ID (Hamada *et al*, 2017) |
| MYO5A | Unconventional myosin | Autism spectrum disorder and synapses (Pandian *et al*, 2020) |
| PAK2 | Serine/threonine-protein kinase | ID (Antonarakis *et al*, 2021) Autism-related behaviors (Wang *et al*, 2018) |
| DCTN1 | Dynactin subunit | Perry syndrome (neurodegeneration) (Dulski *et al*, 1993) |
| LMNA | Prelamin A/C |  |
| SRC | Proto-oncogene tyrosine-protein kinase | Seizures (Liu *et al*, 2022) NMDA receptor and synaptic plasticity (Sinai *et al*, 2010) |
| GNAQ | G(q) protein subunit | Sturge-Weber syndrome with seizures and ID (Day *et al*, 2019) |
| CDK5 | Cyclin-dependent kinase | Neurodegeneration and cognitive dysfunction (Gutiérrez-Vargas *et al*, 2015) ID (Moncini *et al*, 2016) |
| PLCG1 | Phospholipase | Synapse transmission and seizures (Kim *et al*, 2019) |
| MECP2 | Methyl-CpG binding protein | Rett syndrome (Kyle *et al*, 2018) |
| ACTN3 | Alpha-actinin |  |
| HMGB2 | Chromosomal protein | Neural stem cell proliferation and adult neurogenesis (Abraham *et al*, 2013) |
| SYT1 | Membrane-trafficking protein | Mediator of neurotransmitter release, ID, movement disorders (Melland *et al*, 2022) |
| CALM1 | Calcium-binding messenger protein |  |
| ANXA2 | Annexin |  |
| L1CAM | Neural cell adhesion molecule | ID (Bousquet *et al*, 2021; Patzke *et al*, 2016) |
| ANK2 | Ankyrin | Autism (Yang *et al*, 2019; Kawano *et al*, 2022) |
| ACTN2 | Alpha-actinin |  |
| TNNT2 | Tropomyosin-binding subunit of troponin |  |
| MYH6 | Myosin |  |
| ANXA1 | Annexin | Autism spectrum disorder (Correia *et al*, 2014) |
| PTK7 | Tyrosine-protein kinase |  |
| TPM1 | Tropomyosin |  |
| SFRP1 | Secreted frizzled-related protein | Seizures (Diao *et al*, 2021) |
| POSTN | Periostin |  |
| CASQ2 | Calcium-binding protein |  |

**References**

Abraham AB, Bronstein R, Reddy AS, Maletic-Savatic M, Aguirre A & Tsirka SE (2013) Aberrant neural stem cell proliferation and increased adult neurogenesis in mice lacking chromatin protein HMGB2. *PLoS One* 8: e84838

Antonarakis SE, Holoubek A, Rapti M, Rademaker J, Meylan J, Iwaszkiewicz J, Zoete V, Wilson C, Taylor J, Ansar M, *et al* (2021) Dominant monoallelic variant in the PAK2 gene causes Knobloch syndrome type 2. *Hum Mol Genet* 31: 1–9

Arya R, Spaeth C & Zhang W (2021) Epilepsy phenotypes associated with MAP1B-related brain malformations. *Epileptic Disord* 23: 392–396

Bousquet I, Bozon M, Castellani V, Touraine R, Piton A, Gérard B, Guibaud L, Sanlaville D, Edery P, Saugier-Veber P, *et al* (2021) X-linked partial corpus callosum agenesis with mild intellectual disability: identification of a novel L1CAM pathogenic variant. *Neurogenetics* 22: 43–51

Chen X, Sun W, Pan Y, Yang Q, Cao K, Zhang J, Zhang Y, Chen M, Chen F, Huang Y, *et al* (2013) Lithium ameliorates open-field and elevated plus maze behaviors, and brain phospho-glycogen synthase kinase 3-beta expression in fragile X syndrome model mice. *Neurosciences (Riyadh)* 18: 356–362

Cheng Y-Y, Chou Y-T, Lai F-J, Jan M-S, Chang T-H, Jou I-M, Chen P-S, Lo J-Y, Huang S-S, Chang N-S, *et al* (2020) Wwox deficiency leads to neurodevelopmental and degenerative neuropathies and glycogen synthase kinase 3β-mediated epileptic seizure activity in mice. *Acta Neuropathol Commun* 8: 6

Cheon MS, Fountoulakis M, Cairns NJ, Dierssen M, Herkner K & Lubec G (2001) Decreased protein levels of stathmin in adult brains with Down syndrome and Alzheimer’s disease. *J Neural Transm Suppl*: 281–288

Correia CT, Conceição IC, Oliveira B, Coelho J, Sousa I, Sequeira AF, Almeida J, Café C, Duque F, Mouga S, *et al* (2014) Recurrent duplications of the annexin A1 gene (ANXA1) in autism spectrum disorders. *Mol Autism* 5: 28

Day AM, McCulloch CE, Hammill AM, Juhász C, Lo WD, Pinto AL, Miles DK, Fisher BJ, Ball KL, Wilfong AA, *et al* (2019) Physical and Family History Variables Associated With Neurological and Cognitive Development in Sturge-Weber Syndrome. *Pediatr Neurol* 96: 30–36

Diao L, Yu H, Li H, Hu Y, Li M, He Q, Lu L, Li H & Liao X (2021) LncRNA UCA1 alleviates aberrant hippocampal neurogenesis through regulating miR-375/SFRP1-mediated WNT/β-catenin pathway in kainic acid-induced epilepsy. *Acta Biochim Pol* 68: 159–167

Dulski J, Konno T & Wszolek Z (1993) DCTN1-Related Neurodegeneration. In *GeneReviews®*, Adam MP Everman DB Mirzaa GM Pagon RA Wallace SE Bean LJ Gripp KW & Amemiya A (eds) Seattle (WA): University of Washington, Seattle

Fischer B, Retchkiman I, Bauer J, Platt D & Popa-Wagner A (1995) Pentylenetetrazole-induced seizure up-regulates levels of microtubule-associated protein 1B mRNA and protein in the hippocampus of the rat. *J Neurochem* 65: 467–470

Fuchs C, Rimondini R, Viggiano R, Trazzi S, De Franceschi M, Bartesaghi R & Ciani E (2015) Inhibition of GSK3β rescues hippocampal development and learning in a mouse model of CDKL5 disorder. *Neurobiol Dis* 82: 298–310

Fuchs C, Trazzi S, Torricella R, Viggiano R, De Franceschi M, Amendola E, Gross C, Calzà L, Bartesaghi R & Ciani E (2014) Loss of CDKL5 impairs survival and dendritic growth of newborn neurons by altering AKT/GSK-3β signaling. *Neurobiol Dis* 70: 53–68

Guo W, Murthy AC, Zhang L, Johnson EB, Schaller EG, Allan AM & Zhao X (2012) Inhibition of GSK3β improves hippocampus-dependent learning and rescues neurogenesis in a mouse model of fragile X syndrome. *Hum Mol Genet* 21: 681–691

Gutiérrez-Vargas JA, Múnera A & Cardona-Gómez GP (2015) CDK5 knockdown prevents hippocampal degeneration and cognitive dysfunction produced by cerebral ischemia. *J Cereb Blood Flow Metab* 35: 1937–1949

Hamada N, Negishi Y, Mizuno M, Miya F, Hattori A, Okamoto N, Kato M, Tsunoda T, Yamasaki M, Kanemura Y, *et al* (2017) Role of a heterotrimeric G-protein, Gi2, in the corticogenesis: possible involvement in periventricular nodular heterotopia and intellectual disability. *J Neurochem* 140: 82–95

Hertz NT, Adams EL, Weber RA, Shen RJ, O’Rourke MK, Simon DJ, Zebroski H, Olsen O, Morgan CW, Mileur TR, *et al* (2019) Neuronally Enriched RUFY3 Is Required for Caspase-Mediated Axon Degeneration. *Neuron* 103: 412-422.e4

Horn S, Au M, Basel-Salmon L, Bayrak-Toydemir P, Chapin A, Cohen L, Elting MW, Graham JM, Gonzaga-Jauregui C, Konen O, *et al* (2019) De novo variants in PAK1 lead to intellectual disability with macrocephaly and seizures. *Brain* 142: 3351–3359

Jorge-Torres OC, Szczesna K, Roa L, Casal C, Gonzalez-Somermeyer L, Soler M, Velasco CD, Martínez-San Segundo P, Petazzi P, Sáez MA, *et al* (2018) Inhibition of Gsk3b Reduces Nfkb1 Signaling and Rescues Synaptic Activity to Improve the Rett Syndrome Phenotype in Mecp2-Knockout Mice. *Cell Rep* 23: 1665–1677

Kawano S, Baba M, Fukushima H, Miura D, Hashimoto H & Nakazawa T (2022) Autism-associated ANK2 regulates embryonic neurodevelopment. *Biochem Biophys Res Commun* 605: 45–50

Kernohan KD, McBride A, Hartley T, Rojas SK, Care4Rare Canada Consortium, Dyment DA, Boycott KM & Dyack S (2019) p21 protein-activated kinase 1 is associated with severe regressive autism, and epilepsy. *Clin Genet* 96: 449–455

Kim HY, Yang YR, Hwang H, Lee H-E, Jang H-J, Kim J, Yang E, Kim H, Rhim H, Suh P-G, *et al* (2019) Deletion of PLCγ1 in GABAergic neurons increases seizure susceptibility in aged mice. *Sci Rep* 9: 17761

Kyle SM, Vashi N & Justice MJ (2018) Rett syndrome: a neurological disorder with metabolic components. *Open Biol* 8: 170216

Liu L, Xia L, Li Y, Zhang Y, Wang Q, Ding J & Wang X (2022) Inhibiting SRC activity attenuates kainic-acid induced mouse epilepsy via reducing NR2B phosphorylation and full-length NR2B expression. *Epilepsy Res* 185: 106975

Lu R, Wang H, Liang Z, Ku L, O’donnell WT, Li W, Warren ST & Feng Y (2004) The fragile X protein controls microtubule-associated protein 1B translation and microtubule stability in brain neuron development. *Proc Natl Acad Sci U S A* 101: 15201–15206

Luo Y, Shan G, Guo W, Smrt RD, Johnson EB, Li X, Pfeiffer RL, Szulwach KE, Duan R, Barkho BZ, *et al* (2010) Fragile x mental retardation protein regulates proliferation and differentiation of adult neural stem/progenitor cells. *PLoS Genet* 6: e1000898

Melland H, Bumbak F, Kolesnik-Taylor A, Ng-Cordell E, John A, Constantinou P, Joss S, Larsen M, Fagerberg C, Laulund LW, *et al* (2022) Expanding the genotype and phenotype spectrum of SYT1-associated neurodevelopmental disorder. *Genet Med* 24: 880–893

Moncini S, Castronovo P, Murgia A, Russo S, Bedeschi MF, Lunghi M, Selicorni A, Bonati MT, Riva P & Venturin M (2016) Functional characterization of CDK5 and CDK5R1 mutations identified in patients with non-syndromic intellectual disability. *J Hum Genet* 61: 283–293

Ohori S, Mitsuhashi S, Ben-Haim R, Heyman E, Sengoku T, Ogata K & Matsumoto N (2020) A novel PAK1 variant causative of neurodevelopmental disorder with postnatal macrocephaly. *J Hum Genet* 65: 481–485

Pan X, Chang X, Leung C, Zhou Z, Cao F, Xie W & Jia Z (2015) PAK1 regulates cortical development via promoting neuronal migration and progenitor cell proliferation. *Mol Brain* 8: 36

Pandian S, Zhao J-P, Murata Y, Bustos FJ, Tunca C, Almeida RD & Constantine-Paton M (2020) Myosin Va Brain-Specific Mutation Alters Mouse Behavior and Disrupts Hippocampal Synapses. *eNeuro* 7: ENEURO.0284-20.2020

Patzke C, Acuna C, Giam LR, Wernig M & Südhof TC (2016) Conditional deletion of L1CAM in human neurons impairs both axonal and dendritic arborization and action potential generation. *J Exp Med* 213: 499–515

Putra M, Puttachary S, Liu G, Lee G & Thippeswamy T (2020) Fyn-tau Ablation Modifies PTZ-Induced Seizures and Post-seizure Hallmarks of Early Epileptogenesis. *Front Cell Neurosci* 14: 592374

Shaw JL & Chang KT (2013) Nebula/DSCR1 upregulation delays neurodegeneration and protects against APP-induced axonal transport defects by restoring calcineurin and GSK-3β signaling. *PLoS Genet* 9: e1003792

Shukkur EA, Shimohata A, Akagi T, Yu W, Yamaguchi M, Murayama M, Chui D, Takeuchi T, Amano K, Subramhanya KH, *et al* (2006) Mitochondrial dysfunction and tau hyperphosphorylation in Ts1Cje, a mouse model for Down syndrome. *Hum Mol Genet* 15: 2752–2762

Sinai L, Duffy S & Roder JC (2010) Src inhibition reduces NR2B surface expression and synaptic plasticity in the amygdala. *Learn Mem* 17: 364–371

Tang X-Y, Xu L, Wang J, Hong Y, Wang Y, Zhu Q, Wang D, Zhang X-Y, Liu C-Y, Fang K-H, *et al* (2021) DSCAM/PAK1 pathway suppression reverses neurogenesis deficits in iPSC-derived cerebral organoids from patients with Down syndrome. *J Clin Invest* 131: e135763, 135763

Trazzi S, Fuchs C, De Franceschi M, Mitrugno VM, Bartesaghi R & Ciani E (2014) APP-dependent alteration of GSK3β activity impairs neurogenesis in the Ts65Dn mouse model of Down syndrome. *Neurobiol Dis* 67: 24–36

Walters GB, Gustafsson O, Sveinbjornsson G, Eiriksdottir VK, Agustsdottir AB, Jonsdottir GA, Steinberg S, Gunnarsson AF, Magnusson MI, Unnsteinsdottir U, *et al* (2018) MAP1B mutations cause intellectual disability and extensive white matter deficit. *Nat Commun* 9: 3456

Wang Y, Zeng C, Li J, Zhou Z, Ju X, Xia S, Li Y, Liu A, Teng H, Zhang K, *et al* (2018) PAK2 Haploinsufficiency Results in Synaptic Cytoskeleton Impairment and Autism-Related Behavior. *Cell Rep* 24: 2029–2041

Xu L, Huo H-Q, Lu K-Q, Tang X-Y, Hong Y, Han X, Fu Z-X, Fang K-H, Xu M, Guo X, *et al* (2022) Abnormal mitochondria in Down syndrome iPSC-derived GABAergic interneurons and organoids. *Biochim Biophys Acta Mol Basis Dis* 1868: 166388

Yang R, Walder-Christensen KK, Kim N, Wu D, Lorenzo DN, Badea A, Jiang Y-H, Yin HH, Wetsel WC & Bennett V (2019) ANK2 autism mutation targeting giant ankyrin-B promotes axon branching and ectopic connectivity. *Proc Natl Acad Sci U S A* 116: 15262–15271

Zhao F, Hu Y, Zhang Y, Zhu Q, Zhang X, Luo J, Xu Y & Wang X (2012) Abnormal expression of stathmin 1 in brain tissue of patients with intractable temporal lobe epilepsy and a rat model. *Synapse* 66: 781–791
